# An all-in-one AAV vector for cardiac-specific gene silencing by an adenine base editor

**DOI:** 10.1101/2024.09.30.615742

**Authors:** Zhanzhao Liu, Luzi Yang, Yuhan Yang, Jiting Li, Zhan Chen, Congting Guo, Qianhao Guo, Qiuxuan Li, Yueshen Sun, Dongyu Zhao, Xiaomin Hu, Fei Gao, Yuxuan Guo

## Abstract

ABE editing outcomes are highly variable and unpredictable, depending on various factors. Therefore, the success rate of creating targeted A•T-to-G•C conversions using ABE is not high. But utilizing ABE to target RNA splicing sites for gene silencing has a higher success rate. Another challenge for base editing in the heart is that traditional ABE is too large in size, necessitating dual AAV delivery. Whether single AAV delivery can be achieved remains to be explored. In this study, we demonstrated how the diversity of Cas9 homologs and screening of sgRNAs can facilitate cardiac base editing. Single-AAV base editing outperformed dual-AAV systems in the heart, potentially benefiting from chromatin accessibility-guided sgRNA selection. These findings have important implications for the development of more effective and predictable base editing tools for cardiac gene therapy.

## Main text

Adenine base editors (ABEs) are powerful tools to create targeted A•T-to-G•C conversions. An array of studies applied ABEs to correct genetic variants^1^ or ablate post-translational modifications^2^, holding a tremendous potential for cardiac gene therapy. However, the editing outcomes of ABEs are highly variable and unpredictable, depending on the editor properties, target sequence contexts and chromatin accessibility^3^. A large fraction of adenines could not be edited by ABE at an acceptable efficiency, leaving a major technical challenge in ABE applications. Potential solutions include the adaptation of diverse editors with distinct properties and protospacer adjacent motif (PAM) ^4^. In addition, the application of ABE in gene silencing is more likely to succeed than in targeted amino-acid switching because the former strategy could screen more sgRNAs targeting start codon or splice sites for the one with best activity. However, these strategies are yet to be systemically applied to the heart.

Another challenge in cardiac base editing is the large size of canonical ABEs, which deposit limitations in gene delivery. Split-intein-based dual-AAV vectors are currently the major vehicles for cardiac ABE delivery^1, 2^. Several miniature ABE variants that could fit into single AAV vectors have been reported^4^, but how these new ABEs could be efficiently harnessed to edit the heart remain poorly characterized. Whether all-in-one AAV vectors outperform the dual-AAV vectors lacks evidence support.

To solve these problems, we adopted a tool kit of miniature ABEs including SaKKH-ABE, Nme2-ABE, Sauri-ABE and Cj-ABE^4^ with Cas9 features shown in Figure A(1). Eighteen sgRNAs were designed to target A•T in the start codon or 5’ splice sites (5’SS) of *Camk2d* coding CaMKIIδ, a classic therapeutic target for heart diseases^2^. Amplicon sequencing (Amp-seq) screening in N2a cells identified sgRNA1, 6 and 7 of SauriABE as potent sgRNAs for further investigation. Their target sequences are highly conserved and the high activity of their sgRNA homologs in human are validated in HEK293T cells (Figure A(2)).

**Figure.**
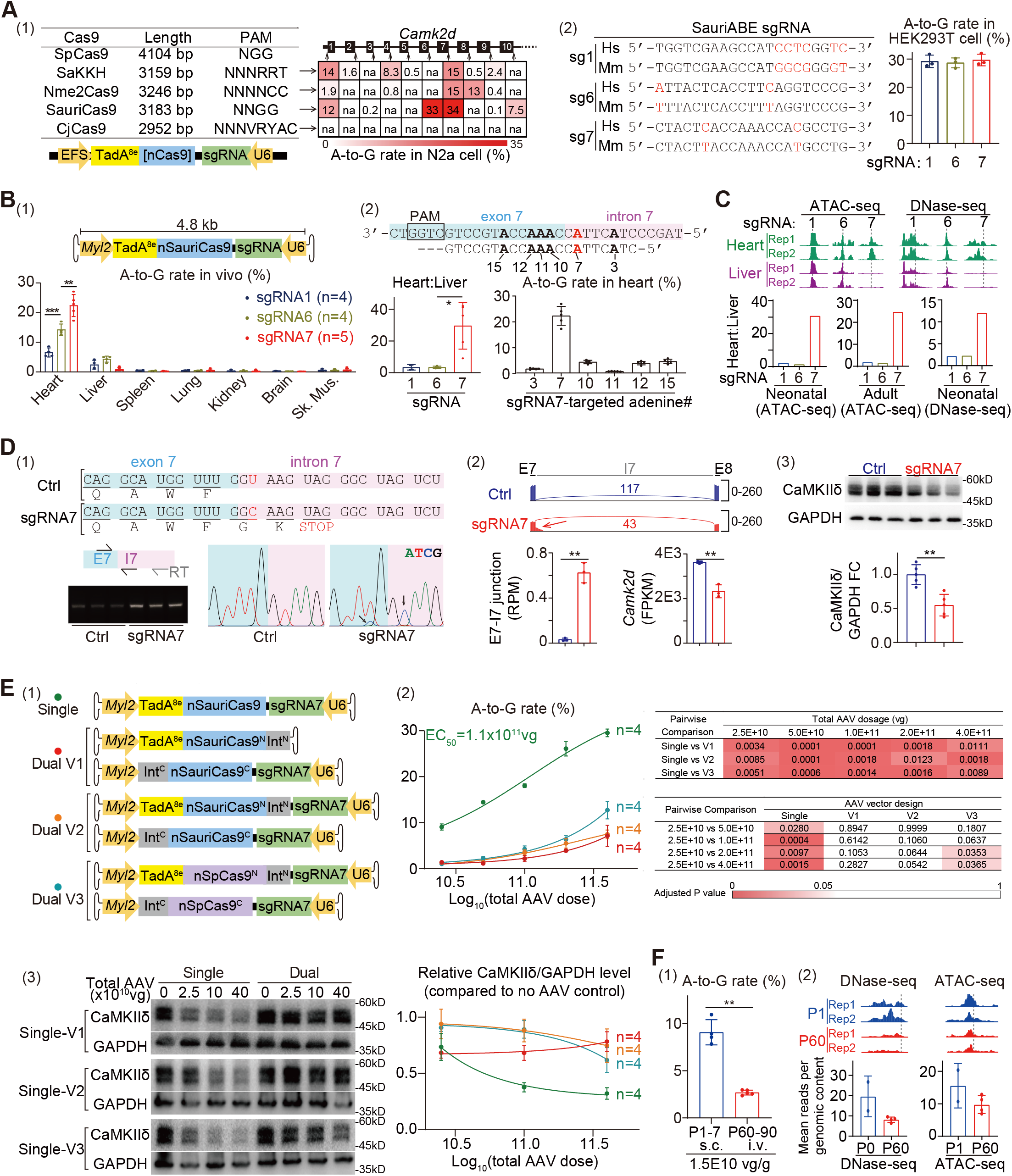
Heart-specific ABE-mediated gene silencing of *Camk2d* via a single AAV vector. **A**, (1) Information of miniature Cas9 variants and sgRNA screen results in murine N2a cells. Plasmids were transfected for 48h before genome DNA was extracted for Amp-seq analysis by CRISPResso2. N = 6 biological repeats. Average editing rates in heatmap. NA, no available sgRNA due to PAM limitation. (2) sgRNA alignments between mouse (Mm) and human (Hs) with mismatches in red and base editing results in human HEK293T cells. Mean ± SD. **B**, (1) AAV-SauriABE editing in various organs. Sk. Mus., skeletal muscle. (2) Base editing ratios and detailed editing results by sgRNA7. P1 mice were treated with 1× 10^11^vg AAV and tissues were collected at P7 for Amp-seq. N = 4-5 mice. Mean ± SD. Unpaired t test with Welch’s correction: *p<0.05, **p<0.01, ***p<0.001. **C**, ATAC-seq and DNase-seq peaks illustrated by integrative genome viewer and peak intensity (±200bp surrounding the sgRNA target site) ratios between heart and liver. Average signal intensities were quantified using pyBigWig. Neonatal ATAC-seq (PRJNA63471, N = 2), adult ATAC-seq (PRJNA497970, N = 4) and neonatal DNase-seq (PRJNA63471, N = 2) were available at NCBI websites. **D**, (1) ABE/sgRNA7 disrupts the 5’SS in exon 7 (E7)-intron 7 (I7) junction and deposits a premature stop codon validated by RT-PCR and sanger sequencing. Arrows point to two cytosine signals due to ABE. (2) Sashimi plots of cardiac RNA-seq data with the numbers of reads containing the E7-E8 junction. The arrow points to aberrant reads covering the E7-I7 junction. RPM, reads per million. FPKM, fragments per kilobases per million. (3) Western blot validation. FC, fold change. Ctrl, control hearts with no AAV treatment. Mean ± SD. Unpaired t test with Welch’s correction: **p<0.01. **E**, (1) the design of single- and dual-AAV vectors. (2) the dose-effect relationship. AAV was injected at P1 and Amp-seq was performed at P7. In dual AAV groups, two AAVs were administered at an equal amount. Mixed-effect ANOVA with Tukey’s multiple comparison test with Padj shown in heatmap. (3) Western blot analysis and quantification. Mean ± SE. **F**, (1) Base editing in neonates versus adults. vg/g, vector genome count per gram body weight. s.c., subcutaneous injection. i.v., intravenous injection. Unpaired t test with Welch’s correction: **p<0.01. (2) DNase-seq and ATAC-seq signal intensities in neonatal versus adult hearts. Mean ± SD. Neonatal and adult DNase-seq data were from PRJNA63471. Neonatal ATAC-seq data were from PRJNA63471, adult data from PRJNA497970.

The *Tnnt2* promoter is commonly used in cardiac specific AAV gene expression, but its large size (>400bp) and leaky liver activity prompted us to test other promoters such as the *Myl2* promoter (<300bp)^5^. We subcutaneously injected 1×10^11^vg (vector genome) AAV9 into postnatal day 1 (P1) C57BL/6 mice and collected ventricles at P7 for Amp-seq analysis. SgRNA7 exhibited the highest editing rate and the best tissue specificity (Figure B(1-2)). Although the exon7-intron7 junction is adenine-rich, sgRNA7 preferentially edited the target adenine in 5’SS over the other bystander adenines (Figure B(2)).

The tissue specificity of AAV-ABE was quantified by heart-to-liver editing ratio, which showed that sgRNA7 intrinsically exhibited cardiac specificity over other sgRNAs. Because chromatin accessibility is a key factor determining genome editing outcomes^3^, three publicly available datasets of ATAC-seq or DNase-seq were analyzed to compare sgRNA-targeted loci (NCBI BioProject PRJNA63471 and PRJNA497970). Strikingly, the sgRNA7-targeted locus was localized in an open region selectively in the heart but not liver, implying tissue-specific chromatin accessibility as a key determinant of organ-selective sgRNA activity (Figure C).

The editing outcome of SauriABE/sgRNA7 was next assessed by reverse-transcription-PCR (RT-PCR), which showed an increase of mRNA containing intron7 (Figure D(1)). Sanger sequencing confirmed the robust editing of intron7 5’SS (Figure D(1)). RNA-seq further demonstrated the prominent increase of mRNA retaining intron7 in the SauriABE/sgRNA7-treated heart while total *Camk2d* mRNA significantly decreased, which is an expected consequence of premature stop codon deposition followed by non-sense-mediated mRNA decay (Figure D(2)). Western blot further validated CaMKIIδ depletion by this AAV-ABE vector (Figure D(3)).

Next, we constructed three dual-AAV systems (Figure E(1)) that carried SauriABE with one copy of U6-sgRNA7 (V1), SauriABE with two copies of U6-sgRNA7 (V2), or SpCas9-ABE with two copies of U6-sgRNA7 (V3). In V1 and V2, SauriABE was split at the site homologous to the conventional dual-ABE vectors like V3. Because SauriCas9 and SpCas9 share almost identical PAM sequences (Figure A(1)), sgRNA7 is almost the same among V1, V2 and V3. The single-AAV vector was side-by-side compared to these three dual-AAV systems by profiling a dose-editing relationship. Strikingly, at all doses, the single-AAV vector triggered cardiac base editing more efficiently than the dual-AAV systems (Figure E(2)), which was further validated by western blot analysis (Figure E(3)). The single-AAV vector triggered higher editing rates at the neonate stage than at the adult stage (Figure F(1)), which correlated with the lower chromatin accessibility in adult hearts (Figure F(2)).

In summary, this study showed how the diversity of Cas9 homologs could facilitate cardiac base editing, which was further enhanced by screening a multitude of sgRNAs for gene silencing. Single-AAV base editing outperformed dual-AAV systems in the heart, which could potentially benefit from chromatin accessibility-guided sgRNA selection.

## Supporting information

Supplemental Figure, and will be used for the link to the file on the preprint site

## Acknowledgements

We thank PackGene Biotech for AAV production and GenePlus for next-generation sequencing.

## Author Contribution

Y.G. conceived and supervised the study. Z.L. designed and constructed the AAV-ABE vectors. L.Y. managed the animal studies. Y.Y. and D.Z. analyzed ATAC-seq and DNase-seq data. Z.C. and Z.L. performed bioinformatic analysis. Z.L. and J.L. performed sgRNA screen in cell culture. C.G., Y.S. and X.H. assisted in animal studies. Q.G., Q.L. and F.G. were responsible for AAV production and characterization. Y.G., Z.L. and Y.Y. prepared the manuscript.

