## Supplemental Figure, and will be used for the link to the file on the preprint site for "An all-in-one AAV vector for cardiac-specific gene silencing by an adenine base editor"

### Uncropped western blot images for Figure D (N=5 animals per group)

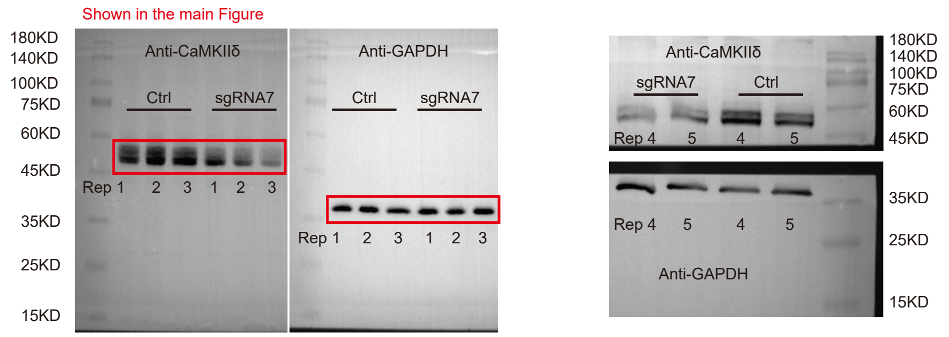


### Uncropped western blot images for Figure E (N=4 animals per group)

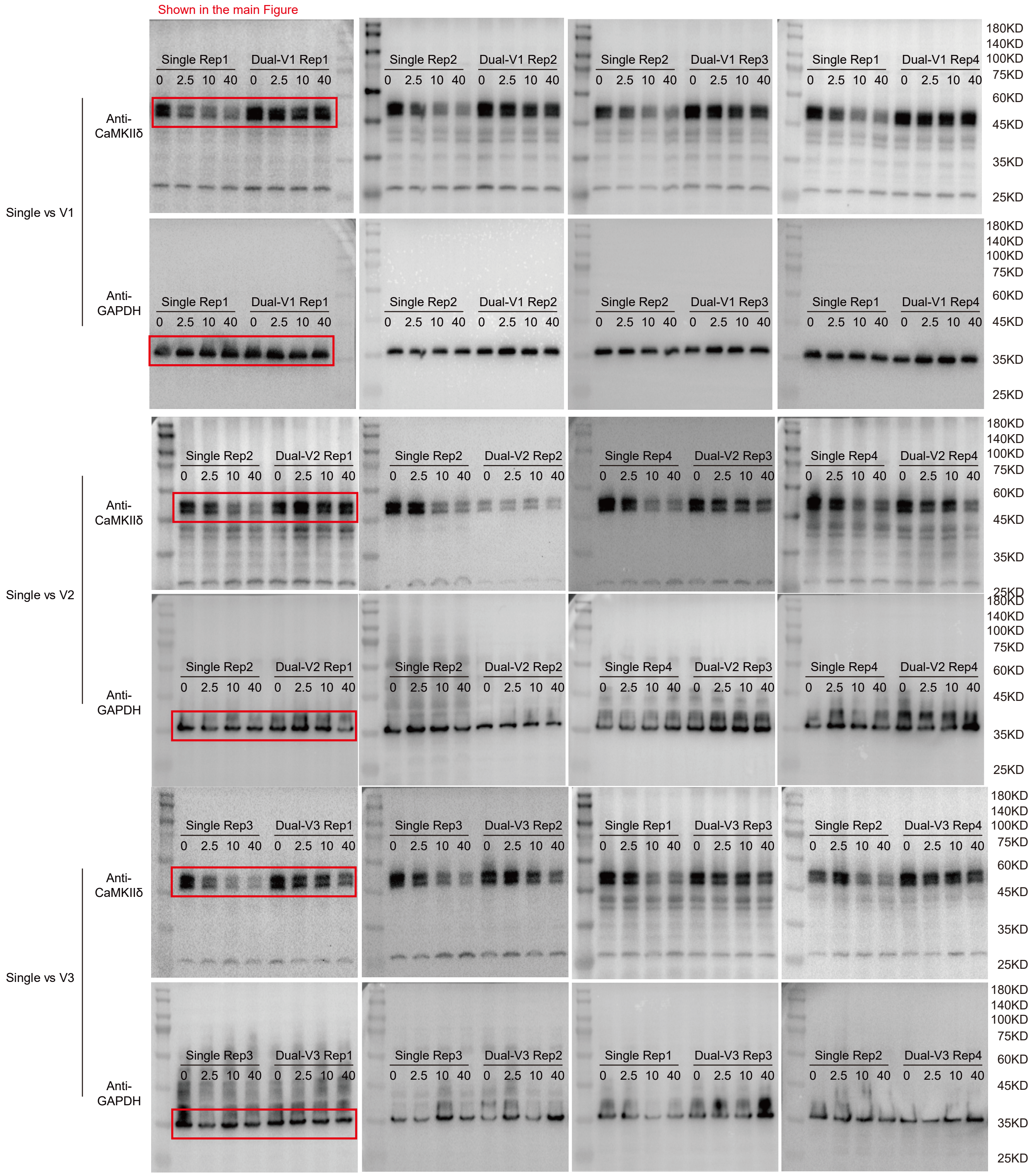
